# 4D live-cell imaging of microgametogenesis in the human malaria parasite *Plasmodium falciparum*

**DOI:** 10.1101/2021.07.28.454129

**Authors:** Sabrina Yahiya, Sarah Jordan, Holly X. Smith, David C. A. Gaboriau, Mufuliat T. Famodimu, Farah A. Dahalan, Alisje Churchyard, George W. Ashdown, Jake Baum

**Affiliations:** Department of Life Sciences, Imperial College London, Sir Alexander Fleming Building, Exhibition Road, South Kensington, London, SW7 2AZ, UK; Facility for Imaging by Light Microscopy, Imperial College London, Sir Alexander Fleming Building, Exhibition Road, South Kensington, London, SW7 2AZ, UK; Walter and Eliza Hall Institute of Medical Research, Parkville, Victoria, Australia

**Keywords:** *Plasmodium falciparum*, microgametogenesis, exflagellation, transmission, parasite biology, time-lapse imaging, microscopy, drug discovery, drug phenotype

## Abstract

Formation of gametes in the malaria parasite occurs in the midgut of the mosquito and is critical to onward parasite transmission. Transformation of the male gametocyte into microgametes, called microgametogenesis, is an explosive cellular event and one of the fastest eukaryotic DNA replication events known. The transformation of one microgametocyte into eight flagellated microgametes requires reorganisation of the parasite cytoskeleton, replication of the 22.9 Mb genome, axoneme formation and host erythrocyte egress, all of which occur simultaneously in <20 minutes. Whilst high-resolution imaging has been a powerful tool for defining stages of microgametogenesis, it has largely been limited to fixed parasite samples, given the speed of the process and parasite photosensitivity. Here, we have developed a live-cell fluorescence imaging workflow that captures the explosive dynamics of microgametogenesis in full. Using the most virulent human malaria parasite, *Plasmodium falciparum*, our live-cell approach combines three-dimensional imaging through time (4D imaging) and covers early microgametocyte development through to microgamete release. Combining live-cell stains for DNA, tubulin and the host erythrocyte membrane, 4D imaging enables definition of the positioning of newly replicated and segregated DNA. It also shows the microtubular cytoskeleton, location of newly formed basal bodies and elongation of axonemes, as well as behaviour of the erythrocyte membrane, including its specific perforation prior to microgamete egress. 4D imaging was additionally undertaken in the presence of known transmission-blocking inhibitors and the untested proteasomal inhibitor bortezomib. Here, for the first time we find that bortezomib inhibition results in a clear block of DNA replication, full axoneme nucleation and elongation. These data not only define a framework for understanding microgametogenesis in general but also suggest that the process is critically dependent on proteasomal activity, helping to identify potentially novel targets for transmission-blocking antimalarial drug development.

## INTRODUCTION

Malaria disease is caused by single-cell protozoan parasites from the genus *Plasmodium*. Over its complex two-host lifecycle, the *Plasmodium* cell demonstrates remarkable cellular plasticity as it transitions between multiple developmental stages. In the transition from mammalian to mosquito host, the parasite faces an extreme population bottleneck in numbers, which also presents a natural target for novel antimalarial treatments aimed at blocking transmission. Transmission is triggered by the uptake of sexual stage gametocytes during a mosquito feed that instantly activate, initiating a transformation in the mosquito midgut that has become a trademark in the cellular biology of these protozoan parasites^1^.

Dormant male (micro) and female (macro) gametocytes form a sub-population of between 0.2-1% of the circulating asexual blood stage parasite reservoir in the mammalian host. The signals that initiate commitment of asexual parasites to sexual differentiation are, however, poorly understood^1^. Committed gametocytes mature over five distinct morphological stages (referred to as stages I-V) and are believed to sequestrate in the host bone marrow and spleen, before emerging into the bloodstream when they reach stage V maturity^1–3^. Following ingestion by a feeding mosquito, stage V microgametocytes and macrogametocytes transform rapidly to microgametes and macrogametes, respectively. The transformation from gametocyte to gamete, a process termed gametogenesis, is activated by a decrease in temperature to 20-25°C, rise in pH and the presence of the mosquito metabolite, xanthurenic acid in the mosquito midgut^4^.

*Plasmodium* gametogenesis is distinctly different between male and female parasites. Both entail a morphological change from falciform to rounded, in *P. falciparum*, and egress from within the host erythrocyte by an ‘inside-out’ mechanism. This mechanism of egress involves disintegration of the parasitophorous vacuole membrane (PVM) prior to that of the host erythrocyte^5^. The female macrogametocyte rounds up^6^ and egresses^7^ within 10 minutes of activation, emerging as a fertilisation competent macrogamete. Whilst this process is incompletely understood, reverse genetic studies using different *Plasmodium* species have described some key female specific events underlying macrogametogenesis^8–12^, such as release of osmiophillic bodies, membrane-bound organelles which are sparse if not entirely absent in microgametocytes^13^. Whilst females become fertilisation-competent upon egress and undergo little cellular reorganisation beyond rounding, microgametogenesis is notoriously complex, and is the focus of our study here.

Microgametogenesis has been most extensively studied by electron microscopy (EM) investigation of the rodent malaria parasite, *P. berghei* and *P. yoelii.^14–16^* The detailed EM work has revealed that microgamete formation entails a stepwise series of events including: substantial cytoskeletal rearrangement, three rounds of DNA replication, alternating with three rounds of endomitotic division, all of which occurs in ~15-20 minutes. *P. falciparum* gametocytes start as falciform, a characteristic from which the species derives its name^2,3^, before morphologically transforming to round once activated. Upon activation, a single microtubule organising centre (MTOC) has been shown to transform into two orthogonal tetrads of basal bodies attached to a spindle pole. The resulting eight basal bodies, from which eight axonemes nucleate and elongate, segregate with each endomitotic division^15,17,18^. Axoneme assembly occurs, fuelled by the large quantities of tubulin within mature microgametocyte that rapidly polymerises to form microtubules^19^. Prior to activation, the MTOC starts in close alignment with the nuclear pore, permitting each basal body to pull a haploid genome (1n) from the newly replicated octoploid (8n) genome through the parental cell body at the point of emergence^15^. This dynamic process by which developing haploid microgametes emerge as motile flagellar is a process termed exflagellation and occurs from ~15 minutes post-activation^17^. Motile haploid microgametes then fuse with the sessile macrogamete, producing a motile zygote able to migrate to the midgut epithelium for oocyst formation and onwards progression in the mosquito^20^.

Current insights into the processes of male and female gametogenesis have taken advantage of the high temporal and spatial resolution offered by brightfield and fluorescence imaging, respectively^21,22^. However, microtubules are not easily resolvable by brightfield and the specificity of fluorescent imaging often requires antibody staining, limiting imaging to fixed samples. This is also true for electron microscopy, despite its proven utility in shaping our current understanding of the fine cellular biology of microgametogenesis^14,15,23^. As a result, the dynamic nature of events encompassing microgametogenesis are still very poorly understood. Better temporal characterisation using live samples, coupled with the specificity of fluorescently tagged structures would allow a marked improvement in our understanding of the process of microgametogenesis and provide a platform from which strategies to block it might then be translatable.

Low and high-resolution microscopy of *Plasmodium* has been extensively used to understand the cell biology of parasite development and aid drug-intervention studies. Ultrastructure expansion microscopy was recently shown to advance traditional fluorescence microscopy approaches to fixed parasite imaging, allowing close observation of asexual blood stage, microgametocyte and ookinete cytoskeletal development^22^. A recent study reported the application of semi-supervised machine learning to define asexual parasite development in a high-throughput imaging format, using fixed parasites^24^. The study proved to be a powerful tool in detecting parasites with morphological perturbations when treated with known antimalarials^24^. Another recent high throughput screen reported the phenotypes of transmission blocking antimalarial hits with unknown cellular targets^25^. The study utilised immunofluorescence labelling of fixed parasites undergoing microgametogenesis to manually define cellular phenotypes. The same screen utilised low-resolution, live brightfield imaging of exflagellation in a high-throughput assay^21^ to identify the microgametogenesis-blocking hits^25^. Live-cell fluorescence microscopy has also been explored, with a recent study utilising lattice light-sheet microscopy to acquire 3D live time-lapse data of asexual *P. falciparum* invasion^26^. Other studies have reported the use of live-cell fluorescence imaging of microgametogenesis to define the phenotypes of transgenic *P. berghei*^27,28^ cell lines. Given the error prone nature of microgametogenesis^16^, however, it is important to define true perturbations to microgametogenesis over natural variation and, critically, to do so in real time.

Here, we describe a protocol that enables the labelling of microtubules, DNA and host erythrocyte membrane for *P. falciparum* microgametocytes and their imaging by live-cell 3D fluorescence microscopy (4D imaging), capturing the entire process of microgametogenesis. To develop a workflow that is translatable to other research labs, we have used a combination of widefield microscopy, an open-source analysis software for deconvolution and commercially available reagents throughout this study. Using this approach, we define in detail the dynamic morphological transformations that occur during microgametogenesis, from activation through to exflagellation. Furthermore, we demonstrate the applicability of our protocol to phenotypic characterisation of inhibitors of microgametogenesis, in particular the role of the proteasome, demonstrating the power of this approach for future transmission-blocking drug discovery^24^.

## RESULTS

### Development of a live microgametogenesis 4D imaging approach

To date, visualisation of the complex cytoskeletal rearrangement, host erythrocyte egress and DNA replication events during *P. falciparum* microgametogenesis (**Figure 1A**) has mostly been limited to fixed imaging protocols. We set out to devise a live cell imaging workflow (**Figure 1B**) that permits observation of cellular dynamics during microgametogenesis in real-time and in three dimensions (4D imaging). Selective testing of several dyes revealed that the silicon-rhodamine (SiR) derivative SiR-tubulin, Vybrant^™^ DyeCycle^™^ Violet and wheat germ agglutinin (WGA) combined to effectively stain live microgametocyte microtubules, DNA and the host erythrocyte membrane, respectively. SiR-tubulin, a non-toxic far-red fluorogenic probe^29^, is an SiR-derivative conjugated to docetaxel^30^ which binds specifically to microtubules and we demonstrate its specificity and photostability in live cell fluorescence imaging of microgametogenesis. Vybrant^™^ DyeCycle^™^ Violet is a cell permeable dye which binds to double-stranded DNA to emit a fluorescent signal proportional to DNA mass and has been previously used to measure microgametocyte genome replication during microgametogenesis^31,32^ using flow cytometry.

**Figure 1.**
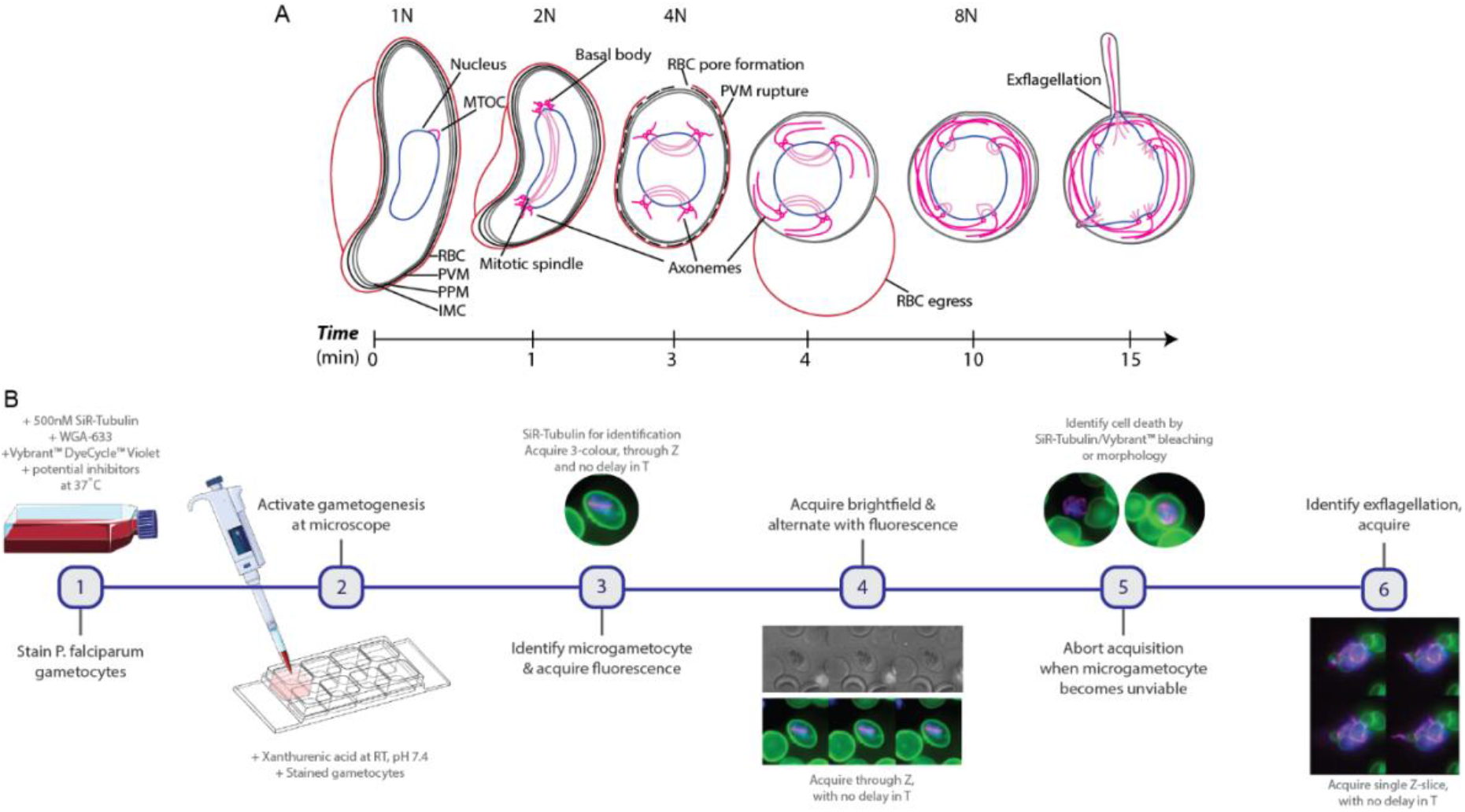
*P. falciparum* microgametogenesis and our live cell imaging approach. **(A)** Details of the cell biological transformations occurring during microgametogenesis from activation at t = 0 minutes to exflagellation at t = 15 minutes. At t = 0 minutes, microgametocytes start with a falciform morphology and an in-tact 4-layer membrane, comprised of the red blood cell (RBC) membrane, parasitophorous vacuole membrane (PVM), parasitophorous plasma membrane (PPM) and inner membrane complex (IMC). Following the first round of DNA replication (1n-2n) at t = 1 minute, the microtubule organising centre (MTOC) transforms to two tetrads of basal bodies joined by a mitotic spindle. Further replication of DNA (2n-4n, 4n-8n) occurs after ~ t = 3-4 minutes, simultaneously to the separation of basal bodies and egress. During egress, PVM rupture precedes erythrocyte egress (inside-out-mechanism) and parasites egress from an erythrocyte pore. Axonemes nucleate from basal bodies following the first DNA replication at t = 1 min and elongate from t = 1-15 min, coiling around the parasite cell body. At 15 minutes post-activation, axonemes emerge attached to a haploid genome as microgametes, in the process of exflagellation. **(B)** The workflow of live gametocyte staining and fluorescence microscopy of microgametogenesis. *P. falciparum* NF54 gametocytes are stained with SiR-Tubulin, WGA-633 and Vybrant^™^ DyeCycle^™^ Violet at 37°C, at which point inhibitors of microgametogenesis may be added. Stained gametocytes are subsequently activated with ookinete medium in pre-positioned imaging slides at RT. SiR-tubulin stained mitotic spindles are used to identify microgametogenesis events, which are continually imaged through T and Z, alternating between brightfield and fluorescence to minimise phototoxicity. When parasites are deemed unviable based on photobleaching or morphology, exflagellation events are captured as single Z-slice timelapses through T.

Stage V gametocytes from the *P. falciparum* NF54 strain, were cultured as previously described^33^, stained and strictly maintained at 37°C to prevent premature activation of gametogenesis. Gametogenesis was initiated by mimicking conditions of the mosquito midgut using “ookinete media” (see Materials and Methods), a xanthurenic-acid-containing media maintained at pH 7.4 and used at room temperature (RT). Labelled gametocytes were directly added to ookinete media-containing imaging slides and prepositioned on the microscope for the immediate acquisition of time-lapse data (**Figure 1B**).

To visualise the initial developmental stages of microgametogenesis, microgametocytes were identified by SiR-tubulin-stained mitotic spindles which signified a successful round of DNA replication (**Figure 1A**). Due to the rapid turnaround between activation and DNA replication, most time-lapses presented here were acquired from 1-2 minutes-post activation, following a round of replication. In optimisation of the imaging workflow, we found the alternation between fluorescence and brightfield acquisition to significantly maximise the viability of microgametocytes. Following identification of an activated microgametocyte, 3-colour fluorescent time-lapses were immediately acquired before switching to brightfield microscopy. A maximum of 10 frames were acquired in fluorescence to minimise the phototoxic effects and brightfield microscopy was subsequently used to monitor parasite development. Upon observation of further parasite differentiation in brightfield, for example by rounding up or preparing for egress, image acquisition was switched back to fluorescence for further acquisition of time-lapses to capture the full early developmental stages of microgametogenesis.

Despite best efforts to reduce LED intensity and frame rates, phototoxicity nearly always prevented the complete visualisation of microgametogenesis from start to finish. To circumvent this issue, activated microgametocytes were imaged at different stages to ensure full capture of later developmental stages, specifically the emergence of microgametes during exflagellation (**Figure 1A**). In this late-stage instance, viable microgametocytes were identified based on SiR-tubulin-stained axonemes, coiled around the parasite cell body. Whilst the earlier stages of microgametogenesis were acquired in 4D, through Z and T, exflagellation could only be captured as single Z-slice time-lapses given the dynamic nature of emerging microgametes (**Figure 1B**). Time-lapse data of the early and later stages of microgametogenesis were subsequently combined and could be analysed together, enabling us to dissect microgametogenesis in its entirety, from initial endomitotic division through to microgamete emergence, for the first time.

### Insights into cytoskeletal rearrangements during microgametogenesis

The formation of mitotic spindles, basal bodies and axonemes occurs with rapid succession during the early stages of microgametogenesis^15,16^. Using our 4D imaging platform we sought to define these stages in real time.

As depicted in **Figures 2A**–**C**, the mitotic spindle of a developing microgametocyte first formed and lengthened across the width of the parasite. Consistent with existing knowledge of microgametogenesis, mitotic spindle formation started out as a single MTOC that then transformed into two tetrads of basal bodies upon the first round of DNA replication (**Figure 1A** and **Supplementary Video 1**). Axonemes were then seen to nucleate from each basal body and subsequently elongated, coiling around the parasite, as shown in **Figures 2A**–**D** and **Supplementary Videos 1-3**. As evident in the 3D data (**Figure 2D** and **Supplementary Video 2**), SiR-tubulin staining gathered at spindle poles whilst four axonemes were nucleated and elongated from basal bodies. We quantified the SiR-tubulin staining intensity of microgametocytes for three distinct developmental stages: microgametocytes with a fully formed spindle; newly nucleated axonemes; and developed axonemes (**Figure 2H**). A significant increase in SiR-tubulin staining was quantified across the three stages (**Figure 2H**), representing the rapid transformation of soluble tubulin into microtubules that occurs during microgametogenesis. This finding was consistent with previous EM studies of microgametogenesis^15,16^. Notably, we have demonstrated the ability to retrieve and quantify volumetric data from our real-time 4D imaging approach which, prior to this study, has not been possible by EM and limited to fixed samples in immunofluorescence studies.

**Figure 2.**
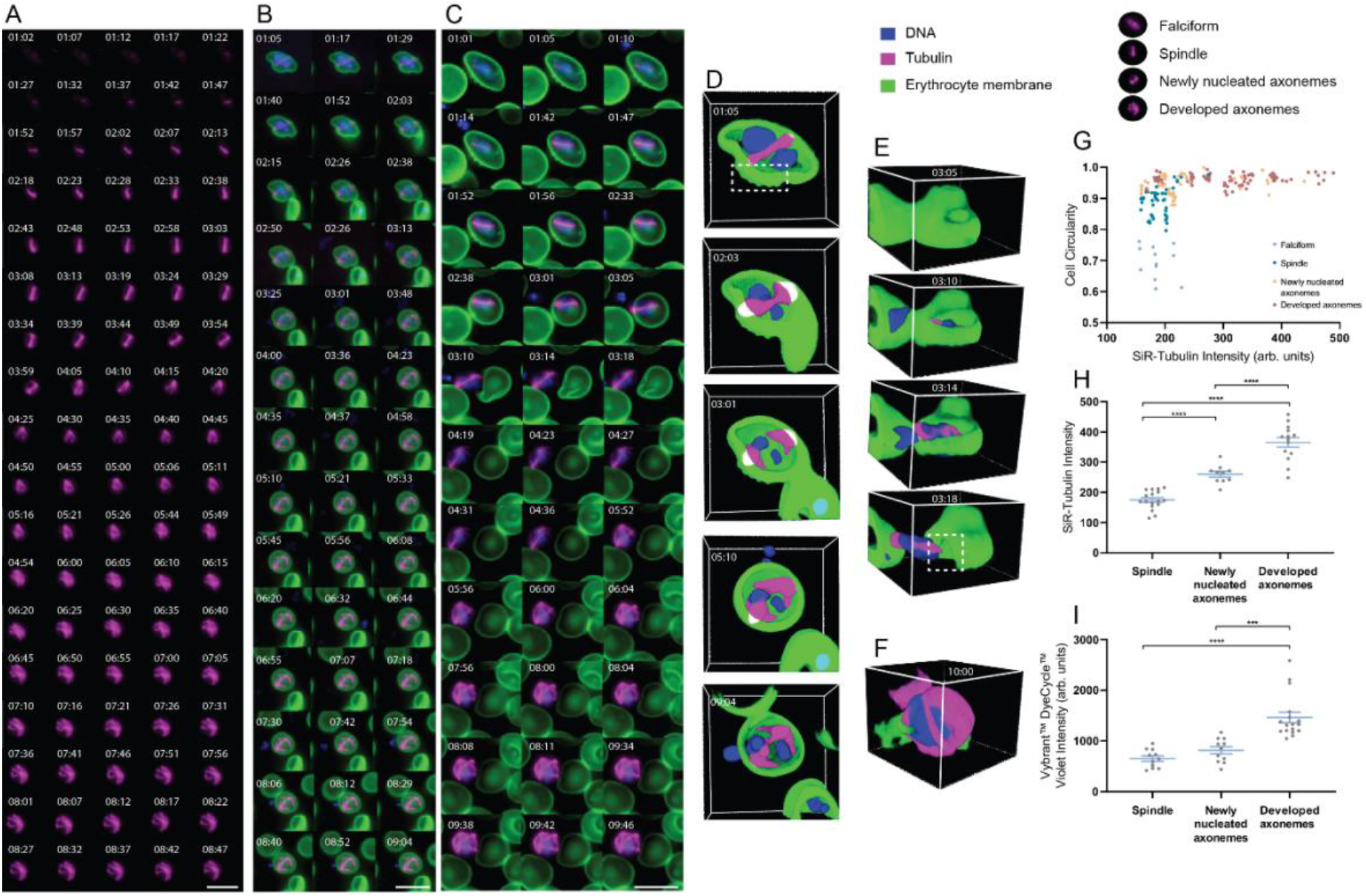
Tubulin dynamics, egress and DNA replication during *P. falciparum* microgametogenesis. Still timelapses of microgametocytes stained with **(A)** SiR-Tubulin (magenta) only and **(B-E)** a combination of SiR-Tubulin (magenta), WGA-488 (green) and Vybrant^™^ DyeCycle^™^ Violet (blue) in the early developmental stages of microgametogenesis. See **Supplementary Videos 1**, **2** and **3** for corresponding timelapses of **A**-**C**. Microtubule staining (magenta) portrays formation of mitotic spindles, basal bodies and axonemes. **(B-E)** Timelapse data depicting DNA replication (blue), microtubule dynamics (magenta), host erythrocyte egress and morphological transformations (host erythrocyte membrane, green). **(D)** Erythrocyte membrane perforation (white dashed box), **(E)** erythrocyte pore formation (white dashed box) and **(F)** axoneme coiling are visible in 3D sectioned images. **(G)** A graph plotted to show cell circularity and SiR-Tubulin intensity (arbitrary units), with each plot representing individual cells of a given developmental stage; falciform (*n* = 12), spindle (*n* = 46), newly nucleated axonemes (*n* = 48) and developed axonemes (*n* = 82). Representative images of developmental stages are above the plots. **(H)** SiR-Tubulin intensity (arbitrary units) of individual cells from distinct developmental stages; spindle (*n* = 17), newly nucleated axonemes (*n* = 10) and developed axonemes (*n* = 14). **(I)** Vybrant^™^ DyeCycle^™^ Violet intensity (arbitrary units) quantified in distinct developmental stages; spindle (*n* = 11), newly nucleated axonemes (*n* = 11) and developed axonemes (*n* = 17). Significant differences in stain intensities between developmental stages was calculated with an unpaired, two-tailed t test (*** *p* < .001, **** *p* < .0001). **A-C** 2D maximum intensity projection of 3D data, scale bars = 10 μm. Individual channels of **B** and **C** can be found in **Figure S1A** and **Figure S1B**, respectively. **D-E** 3D sectioned views frames depicted in **B-C**, respectively. **A-E** Time is depicted as minutes and seconds (mm:ss). All imaging data depicted reflect observations from >10 biological replicates.

Simultaneously to cytoskeletal rearrangement, microgametocyte morphology was seen to transform from falciform to round, transforming the morphology of the host erythrocyte in the same way (**Figure 2B**–**CF** and **Supplementary Videos 2**-**3**). To identify the relationship between rounding up and microtubule polymerisation, we quantified microgametocyte cell circularity and SiR-tubulin staining intensity, respectively. Individual cells were characterised across developmental stages: microgametocytes that were falciform or had a fully developed mitotic spindle, versus those with newly nucleated axonemes or developed axonemes. We found a non-linear relationship between microgametocyte circularity and SiR-tubulin intensity (**Figure 2G**). Individual microgametocytes at the initial developmental stages of microgametogenesis showed varying levels of circularity but a similarly low level of SiR-tubulin intensity. Upon reaching maximal microgametocyte circularity, cells with newly nucleated or developed axonemes showed varying SiR-tubulin intensities, which were at a higher level than the earlier developmental stages. The non-linear relationship between rounding up and microtubule polymerisation revealed that early microtubule polymerisation occurs simultaneously to rounding up and, upon fully rounding up, axonemes continue to elongate and develop (**Figure 2G**)

### Microgametocyte DNA segregates and localises perpendicularly to spindle poles

Incorporating the DNA dye, Vybrant^™^ DyeCycle^™^ Violet, to stain microgametocytes, we next explored the behaviour of the nucleus. Using 3D sectioned data derived from our 4D dataset, we observed nuclear segregation to occur as the microgametocyte genome was replicated (**Figure 2D** and **Supplementary Video 2**). A novel observation was made when comparing the localisation of tubulin and DNA staining, where we observed segregated DNA positioned perpendicularly to basal body tetrads (**Figure 2D**,**Supplementary Video 2** and **Figure 5A**). This showed that the 3D data derived can be used to make novel observations on the biology of microgametocyte DNA replication.

Upon full axoneme development, DNA content visibly increased (**Figure 2F**) compared to earlier stages of microgametogenesis (**Figure 2D**). When quantified, a significant increase of DNA staining from spindle formation to nucleation and development of axonemes was found (**Figure 2I**). This demonstrated that the volumetric data can be obtained using our imaging framework, permitting real-time quantification of DNA content and providing a unique window into genome replication.

### The host erythrocyte perforates and forms a pore for microgametocyte egress

A key stage in microgametogenesis is parasite egress from the host erythrocyte (**Figure 2C, E** and **Supplementary Video 3**). 4D imaging revealed that the erythrocyte membrane perforated in preparation for egress (**Figure 2D** and **5A**). Activated microgametocytes aligned at the periphery of the parasite membrane, eventually ejecting from a frequently singular pore that formed in the host erythrocyte (**Figure 2E** and **Supplementary Video 3**). Notably, we observed microgametocytes eject from a spindle pole of the parasite, out of the newly formed host erythrocyte pore (**Figure 2E**, **Figures S1B-D**, **Figure 5A**, and **Supplementary Videos 3**,**5** and **6**). This finding suggests the mechanism of microgametocyte egress may utilise an unexplored driving force that coordinates ejection from the host cell with spindle pole positioning.

### Real-time fluorescence imaging of exflagellation

The final steps of microgametogenesis are the emergence of haploid microgametes from the parasite cell body, the remarkably dynamic process of exflagellation. Upon full elongation, the tip of an axoneme aligns at the periphery of the parasite cell body to emerge. When emerging at ~15 minutes post-activation, the developed axoneme carries a haploid genome (1n) from the newly replicated octoploid genome (8n) through the cell surface (**Figure 1A**).

Emerging microgametes could be identified either by SiR-tubulin stained axonemes or by the increased motion visible in brightfield, with the latter minimising the effects of phototoxicity. Following the initial emergence of microgametes (**Figure S2** and **Supplementary Video 7**), the full length of axonemes continued to emerge in a rapid motion (**Figure 3**). Full microgamete lengths were visualised by brightfield (**Figure 3A** and **Supplementary Video 8**-**9**) and SiR-tubulin (**Figure 3B, C, E** and **Supplementary Videos 10**-**17**). Host erythrocyte staining during exflagellation revealed the adherence of microgametes to neighbouring erythrocytes (**Figure 3C, E** and **Supplementary Videos 15-17**). DNA staining was notably increased during exflagellation (**Figure 3C, E** and **Supplementary Videos 15-17**) compared to earlier stages of microgametogenesis (**Figure 2B**–**C** and **3**), signifying successful replication.

**Figure 3.**
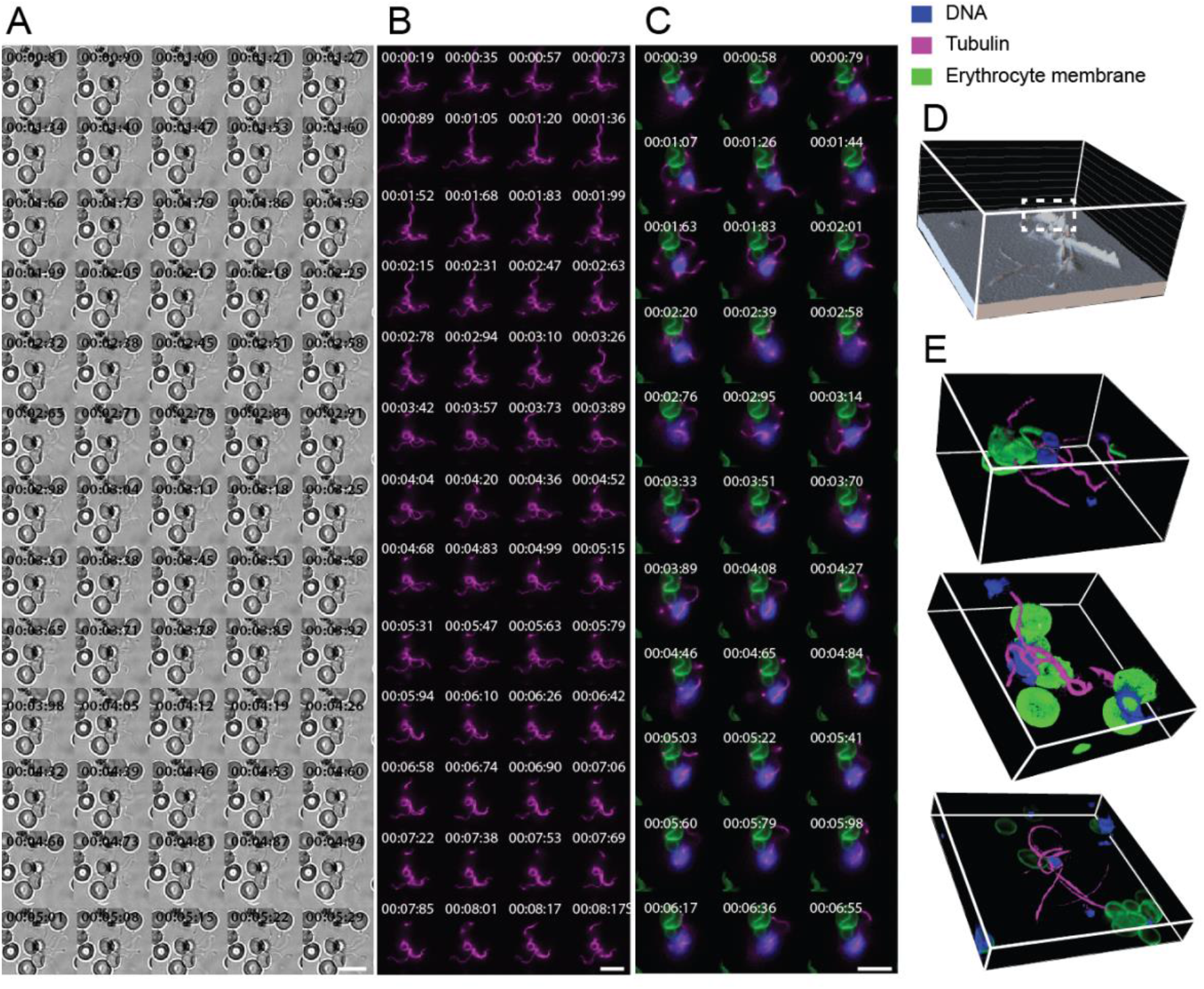
Exflagellation of *P. falciparum* microgametes. 2D timelapse stills of exflagellation imaged by **(A)** brightfield and **(B, C** & **E)** fluorescence microscopy. See **Supplementary Videos 8**, **10** and **15** for corresponding timelapses of **A**-**C**. **(B-C)** SiR-Tubulin-stained axonemes (magenta) emerge from the parasite cell as microgametes. **(C)** Emerging microgametes carry a 1n genome from the newly replicated 8n genome (blue) and adhere to neighbouring erythrocytes (green). **(D)** 3D intensity plot of SiR-Tubulin staining intensity to reveal dense regions (white dashed line) of axoneme overlap. **(E)** 3D-sectioned views of exflagellating microgametes, see **Supplementary Video 17** for a 3D rotated view. **A-C** Time is depicted as minutes, seconds and milliseconds (mm:ss:ms). Scale bars = 10 μm. Individual channels of **C** can be found in **Figure S1E**. All data depicted reflect observations from >10 biological replicates.

Due to the dynamic nature of emerging microgametes that were motile through Z-acquisition, 2D rather than 3D time-lapse data was acquired during exflagellation. Although 3D frames of exflagellation were not obtained, it is possible to obtain 3D plots of SiR-tubulin intensity staining (**Figure 3D**). This allowed the identification of dense regions of SiR-tubulin staining resulting from axoneme overlap, which would otherwise not be deducible without 4D imaging (**Figure 3D**). Alternatively, as parasite motion halts during loss of viability, 3D data can be obtained to closely observe the positioning of emerged microgametes (**Figure 3E** and **Supplementary Video 17**), demonstrating about ability to acquire both 2D and 3D exflagellation time-lapse data.

### Drug inhibition of microgametogenesis

The process of microgametogenesis is tightly synchronised by a series of cell cycle regulators and is consequently sensitive to and the target of known and developmental drug treatments^25^. We sought to apply our 4D imaging approach as a drug discovery tool that could help elucidate the cellular phenotypes of compounds with known and unknown activity against microgametogenesis regulators towards defining their mode or process of action. Compounds 1294^31,34^ and ML10^35^ have been well-established as potent inhibitors of microgametogenesis regulators Ca^2+^-dependent protein kinase 4 (CDPK4) and cyclic-GMP dependent protein kinase G (PKG), respectively. CDPK4 tightly regulates three processes during microgametogenesis: initiation of the first genome replication, mitotic spindle assembly and microgamete motility^31^. PKG has roles in the regulation of Ca^2+^ levels and rounding up during gametogenesis^36,37^. The cellular phenotype of 1294^31,34^ and ML10^35^ have been previously reported using immunofluorescence staining of fixed microgametocytes^38^. Using 4D imaging, we could resolve distinct cellular phenotypes for each drug and its target (**Figure 4A**–**B, D**–**H** and **Supplementary Videos 18**-**24**).

**Figure 4.**
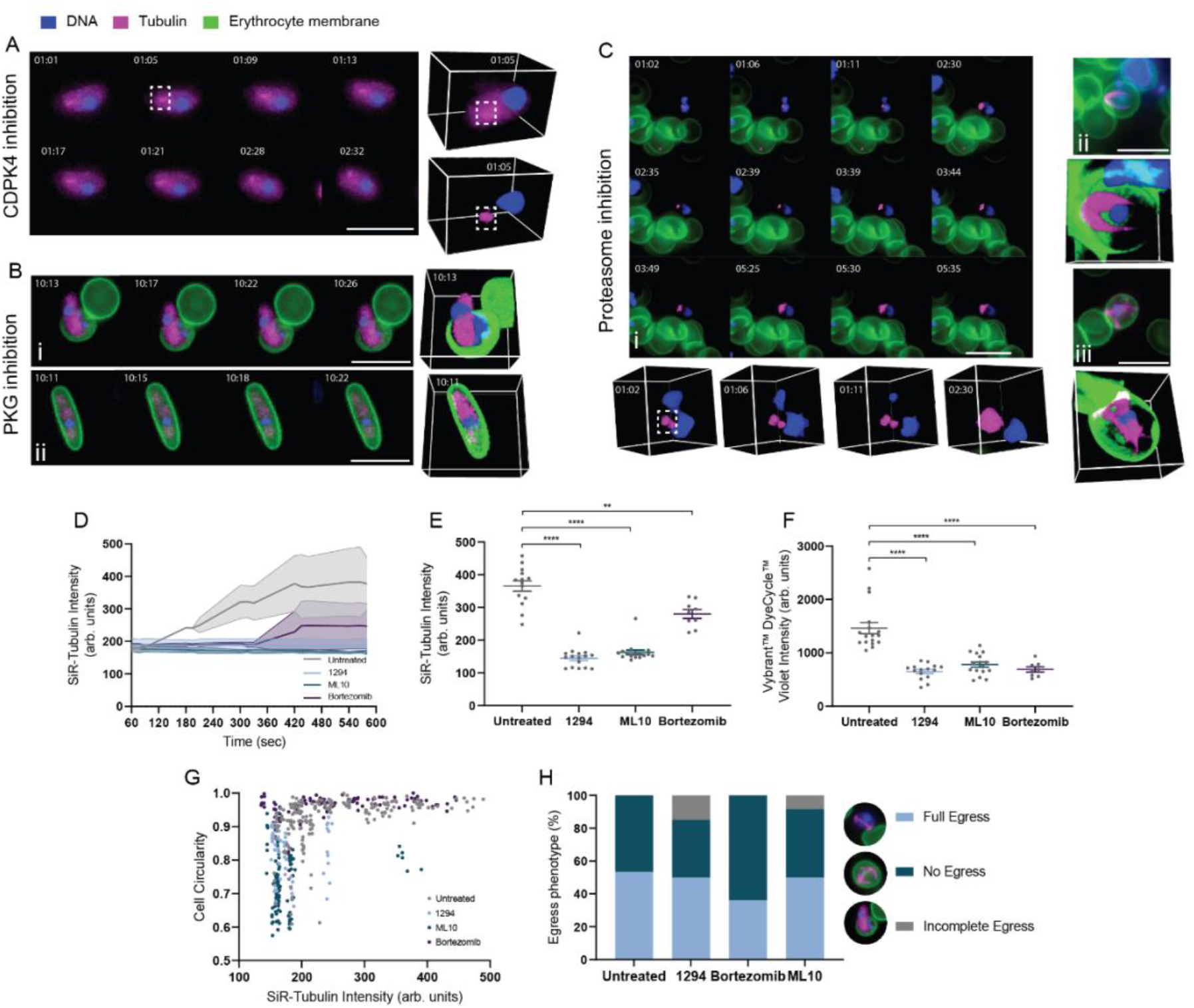
Cellular phenotypes of PKG, CDPK4 and proteasome-inhibited parasites. Cellular phenotypes upon inhibition of *P. falciparum* **(A)** CDPK4, **(B)** PKG and **(C)** proteasome by 1294, ML10 and bortezomib, respectively, during microgametogenesis. Perturbations to microtubule rearrangement (SiR-Tubulin, magenta), the host erythrocyte (WGA, green) and DNA replication (Vybrant^™^ DyeCycle^™^ Violet, blue) are shown as 2D maximum intensity projections of 3D data and alongside 3D sectioned views. Individual channels can be found in **Figure S3**. **(A)** Failed DNA replication, cytoskeletal rearrangement and MTOC (white dashed line) transformation under 1294-treatment are shown. Stress induced egress prior to activation is also depicted. See **Supplementary Video 19** for the corresponding timelapse. **(B)** The failed DNA replication and cytoskeletal rearrangement due to PKG inhibition by ML10 is shown. Mixed egress phenotypes were observed, including **(i)** incomplete and **(ii)** failed egress. See corresponding timelapses in **Supplementary Videos 23-24**. **(C)** Perturbations to **(i)** MTOC transformation (white dashed line) and **(ii-iii)** formation of two-three truncated axonemes resulting from proteasome inhibition are shown. See **Supplementary Videos 25**, **27** and **28** for the corresponding timelapses. **(D)** A continuum of SiR-tubulin staining intensity (arbitrary units) in untreated (*n* = 4), 1294 (*n* = 3), ML10 (*n* = 3) and bortezomib (*n* = 6) treated parasites. **(E)** SiR-tubulin staining (arbitrary units) at 10 minutes post-activation under different treatments. Untreated (*n* = 14), 1294 (*n* = 16), ML10 (*n* = 16), bortezomib (*n* = 9). Significance was calculated with an unpaired, two-tailed t test; ** *p* < .01, **** *p* <.0001. **(F)** Vybrant^™^ DyeCycle^™^ Violet staining (arbitrary units) was significantly reduced (unpaired, two-tailed t test; **** *p* <.0001) at 10 minutes post-activation under different treatments. Untreated (*n* = 17), 1294 (*n* = 15), ML10 (*n* = 9), bortezomib (*n* = 16). **(G)** A graph depicting the cell circularity and SiR-tubulin staining intensity of individual cells across the entirety of microgametogenesis under varying treatments. Untreated (*n* = 188), 1294 (*n* = 58), ML10 (*n* = 106), bortezomib (*n* = 105). **(H)** Percentage egress at 10 minutes post-activation under different treatments was quantified, with distinct egress phenotypes depicted beside the stacked bar graph. Untreated (*n* = 58), 1294 (*n* = 20), ML10 (*n* = 25), bortezomib (*n* = 24). All imaging data depicted reflect observations from >3 biological replicates.

CDPK4 inhibition by 1294^31,34^ prevented morphological transformation from falciform to round (**Figure 4A** and **G**), DNA replication (**Figure 4F**) and microtubule polymerisation (**Figure 4D**–**E**) during microgametogenesis (**Supplementary Videos 18-22**). On detailed inspection, 1294-treated parasites, failed to reach the maximum level of cell circularity (**Figure 4G**) and SiR-tubulin intensity (**Figure 4A, D, E** and **G**), indicating a role of CDPK4 in microgametocyte rounding as well as cytoskeletal rearrangement during microgametogenesis. Of note, many 1294-treated microgametocytes were observed to have egressed from the onset of activation, a probable stress-response of CDPK4 inhibition (**Figure 4A**). Of the population of 1294-treated microgametocytes, 50% were able to fully egress compared to the 53% of untreated microgametocytes which egressed from the host erythrocyte (**Figure 4H**). An incomplete egress phenotype in which falciform parasites partially emerged from the host erythrocyte was also observed, with 15% of 1294-treated parasites demonstrating this phenotype (**Figure 4H**). 3D data revealed the positioning of the MTOC which failed to transform to eight basal bodies (**Figure 4A** and **Supplementary Video 19**). This suggests that CDPK4 plays an early role in DNA replication, microtubule polymerisation, rounding up and host erythrocyte egress during microgametogenesis. This phenotype is consistent with published findings on 1294-treatment of *P. falciparum*^38^ and *P. berghei*^31^ gametocytes, with the exception that our morphological rounding phenotype is not observed in *P. berghei* gametocytes which are already round prior to activation. The deduced overall cellular phenotype of CDPK4 inhibition is summarised in **Figure 5B**.

**Figure 5.**
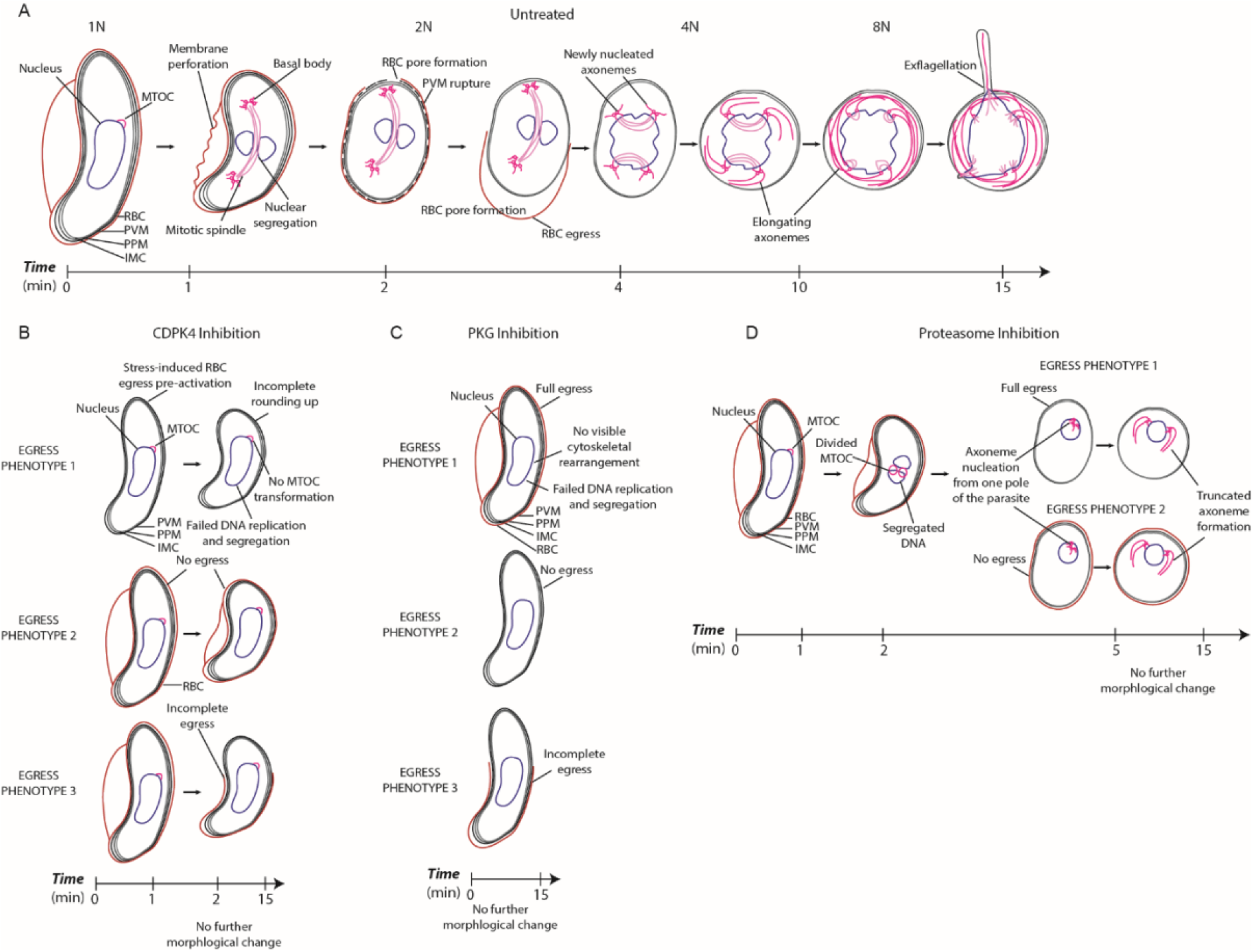
Novel insights into microgametogenesis with and without drug inhibition. Schematic diagrams showing the transformations observed by 4D live-cell fluorescence microscopy of microgametogenesis. The observations depicted are of **(A)** untreated, **(B)** CDPK4-inhibited, **(C)** PKG-inhibited and proteasome-inhibited microgametocytes, treated with DMSO, 1294, ML10 and bortezomib, respectively. **(A)** Perforation and pore-formation of the host cell membrane was found to occur during egress of untreated microgametocytes, which ejected from a single spindle pole of the parasite. Nuclear segregation was found to occur perpendicularly to the mitotic spindle. **(B)** CPDK4 inhibition prevented MTOC transformation and full rounding-up with three distinct egress phenotypes: 1) stress-induced egress prior to activation, 2) no egress and 3) incomplete egress. **(C)** Inhibition of PKG prevented rounding-up and any microtubule polymerisation, with 3 distinct egress phenotypes: 1) no egress, 2) full egress and 3) incomplete egress. **(D)** Proteasome inhibition resulted in abhorrent MTOC division and some nuclear segregation, with few truncated axonemes nucleating from one pole of the transformed parasite. Rounding-up of proteasome-inhibited parasites was observed.

PKG inhibition by ML10^35^ was observed next, clearly demonstrating arrest of microgametocytes before cell-rounding (**Figure 4B** and **G** and **Supplementary Videos 23**-**24**), replication of DNA (**Figure 4F**) and microtubule polymerisation (**Figure 4D** and **E**). ML10-treated gametocytes retained a falciform morphology and a level of SiR-tubulin staining on par with that seen at the onset of gametogenesis activation (**Figure 4D**–**E** and **G** and **Supplementary Videos 23**-**24**). SiR-tubulin intensity was significantly lower than untreated microgametocytes at 10 minutes post-activation (**Figure 4E**). The host erythrocyte egress phenotype was mixed, as 50% of ML10-treated cells emerged fully, 50% failed (**Figure 4Bii**) and 8% partially emerged (**Figure 4Bi**) from the host erythrocyte (**Figure 4A, H** and **Supplementary Videos 23**-**24**). We can deduct from these images that PKG plays a significant role in regulating MTOC transformation, axoneme nucleation and elongation, DNA replication and rounding up exhibited in microgametogenesis, as summarised in **Figure 5C**. These observations match previously reported studies on PKG during microgametogenesis^38,39^ and does so without the laborious staining steps of fixed parasite imaging.

### The microgametocyte proteasome plays a crucial role in microgametogenesis

To extend the cellular dissection of microgametogenesis, we next sought to test the role of the proteosome in cellular reorganisation of the microgametocyte using the drug bortezomib, a proteasome-inhibitor that has not been explored during microgametogenesis. Bortezomib is active against the asexual blood stages of *Plasmodium*^40^ and the eukaryote and euryarchaeota proteasomes^41–43^. A recent study on the 20S proteasome of *S. acidocaldarius* reported bortezomib arrested cells in the midst of division^44^. Given the role of the proteasome in *S. acidocaldarius* cell division and interest in its use as an antimalarial drug target^45^ we aimed to deduce the role of the proteasome in regulation of microgametogenesis DNA replication and cytoskeletal rearrangement.

Proteasome inhibition by bortezomib resulted in a block of microgametogenesis, preventing full cytoskeletal rearrangement (**Figure 4D, E** and **G**), DNA replication (**Figure 4F**) and exflagellation (**Figure 4C**). Most cells were also shown to fail to egress from the host erythrocyte, although this phenotype was mixed (**Figure 4H**). We observed inhibition of the *P. falciparum* proteasome impacted the transformation of the microgametocyte MTOC, visible as 2 small nodes of SiR-tubulin staining which remained at one pole of the parasite (**Figure 4Ci** and **Supplementary Videos 25**). The transformed MTOC was subsequently able to nucleate axonemes from one end of the parasite, but no more than 3 axonemes were formed and growth was truncated (**Figure 4Cii-iii** and **Supplementary Videos 25**-**28**). This perturbation to axoneme nucleation and elongation resulted in a significant decrease in SiR-tubulin staining intensity (**Figure 4D, E** and **G**), although less significant than the decrease observed with ML10 and 1294 treatment which blocked all microtubule polymerisation. Proteasome-inhibited microgametocytes were, however, able to transform from falciform to round (**Figure 4G** and **Supplementary Videos 25**-**28**).

Bortezomib treatment also significantly reduced Vybrant^™^ DyeCycle^™^ Violet staining intensity, signifying a probable indirect inhibitory effect on DNA replication (**Figure 4F**). Additionally, we observed incomplete transformation of the MTOC which resulted in truncated formation of few axonemes from one pole of developing microgametocytes. Combined these data represent the first time that the cellular role of the proteasome during microgametogenesis has been explored. Our findings, summarised in **Figure 5C**, add further weight to bortezomib’s use as a desirable antimalarial drug candidate that is able to inhibit the sexual stages of *Plasmodium* in additional to asexual replication, representing the potential to treat symptoms and block transmission with a single compound.

## DISCUSSION

To date, detailed observation of *P. falciparum* microgametogenesis has mostly centred around fixed parasite imaging, with studies utilising immunofluorescence labelling^25^ or electron microscopy^14–16^. Although these studies have been pivotal in developing our current understanding of microgametogenesis cell biology, imaging fixed parasites is limited by the extensive sample preparation steps and fails to resolve the dynamic nature of underlying cellular events. Here, we have developed a live-cell 3D fluorescence imaging approach (4D imaging) that captures the dynamics of *P. falciparum* microgametogenesis from activation to exflagellation in fine detail. Utilising widefield microscopy, commercially available stains, *in vitro P. falciparum* culture and an open-source software for analysis we present a methodological approach that will be readily utilisable for other research groups.

Using a combination of a cell-permeable fluorogenic probe, DNA dye and lectin we can label and observe development of microgametocyte microtubules, DNA and the host erythrocyte membrane, respectively. Whilst live fluorescence imaging is often impeded by phototoxicity, here we have devised a method that maximises the length of time-lapse image acquisition without compromising microgametocyte viability. Our approach permits the acquisition of microgametogenesis in full over two stages: early and late development, capturing spindle formation through to full axoneme development and exflagellation, respectively. Importantly, our imaging approach permits volumetric quantification of 3D data through time which we demonstrate as a powerful tool in defining drug phenotypes. Our approach is consequently applicable to future comparative studies of alternative drug treatment and *P. falciparum* transgenic cell lines to wild-type phenotypes.

In line with previous studies on microgametogenesis^13,16^, we observed the rapid production of basal bodies from a single MTOC to initiate nucleation and elongation of axonemes, simultaneously to DNA replication and egress. Coupling live fluorescence microscopy to fluorescence intensity and 3D analyses, we observed nuclei to segregate and align perpendicularly to basal bodies in the early stages of microgametogenesis. Whilst synchronous segregation of developing microgametocyte genomes and basal bodies has been reported^13^, live fluorescence imaging provides novel insight into the positioning of the newly replicated DNA. Furthermore, we find egress involves perforation and pore formation of the host erythrocyte membrane. Previous studies have reported swelling of the host erythrocyte prior to PVM rupture and vesiculation during *P. berghei* microgametogenesis^7^. Here, upon erythrocyte pore formation we observed egress of developing *P. falciparum* microgametocytes to occur from a spindle pole of the microgametocyte. This suggests that there may be forces occurring from a single pole of the microgametocyte that drive egress from the host erythrocyte. Future investigation of pore-forming proteins and their role in this process may also be of relevance.

Additionally, we demonstrate the applicability of our workflow to the study of transmission blocking drug phenotypes. We have used the live imaging framework to elucidate the cellular phenotypes of 1294 and ML10, known inhibitors of microgametogenesis regulators CDPK4 and PKG, respectively. We have also defined the proteasome as a crucial component of microgametogenesis regulation and for the first time, we define bortezomib as a powerful inhibitor of *P. falciparum* transmission. This finding points to the importance of the degradation of misfolded proteins and regulation of functional protein abundance in permitting transmission of the *Plasmodium* microgametocytes. Our approach generates reproducible and consistent phenotypes to fixed parasite studies whilst, critically, not requiring complex fixation or staining steps.

As drug and insecticide resistance has threatened existing antimalarial treatment strategies, there is an urgent need for novel transmission-blocking antimalarials that can be used singularly or in combination with schizonticides, killing asexual stages. Developing a full understanding of the mode of action of antimalarial drug candidates maximises the likelihood of clinical safety and future administration. Our imaging approach permits deeper understanding of this remarkable cell biology process, capturing real-time development with fluorescence which may otherwise be missed with fixed or brightfield imaging. The data depicted here promises to unveil novel insights into *P. falciparum* microgametogenesis for cell biology and drug study, but the protocol is not limited to this. Cultivation of *in vitro Plasmodium* cultures, at any stage, permits the live microscopy of the breadth of malaria parasite development, from macrogametogenesis and asexual blood stage development to liver stages. Additional stains for intracellular organelles, parasite membranes and sex-specific proteins, with both wild-type and transgenic lines, will now be a priority for exploring so that we can shed further light on this ancient but deadly single-celled parasite.

## MATERIALS AND METHODS

### In vitro culture of Plasmodium falciparum

*P. falciparum* NF54 strain parasites were cultured as previously described^33^. Asexual parasite cultures were maintained between 0.75-5% parasitaemia and 4% haematocrit using human erythrocytes (NHS National Blood Service). Erythrocytes were supplemented with 3 units/ml heparin (Sigma-Aldrich). Parasites were grown in asexual parasite culture medium (RPMI 1640 with 25 mM HEPES (Life Technologies) supplemented with 50 μg/ml hypoxanthine (Sigma), 0.3 g/l L-glutamine (Sigma) and 10% human serum (Interstate Blood-Bank)). Gametocyte cultures were induced from asexual parasite cultures at 3% asexual parasitaemia and 4% haematocrit. Gametocyte culture media (RPMI 1640 with 25 mM HEPES supplemented with 150 μg/ml L-glutamine, 2.78 mg/ml sodium bicarbonate, 2 mg/ml D-glucose, 50 μg/ml hypoxanthine, 5% human serum and 5% AlbuMAX-II (Gibco)) was replaced daily until reaching maturity at day 14-post induction. All cultures were maintained at 37°C under 3% O_2_/5% CO_2_/93% N_2_ (BOC, UK).

Upon reaching maturity, gametocyte viability was determined by measuring the rate of exflagellation relative to erythrocyte density. Gametocyte culture was treated with ookinete media (RPMI 1640 supplemented with 2 g/l sodium bicarbonate, 50 mg/l hypoxanthine and 100 mM xantharinic acid (XA) (Sigma-Aldrich), pH adjusted to 7.4) to activate gametogenesis. Exflagellation events and erythrocyte density was counted using a haemocytometer (VWR) and Nikon Leica DC500 microscope.

### Staining and Treating Live P. falciparum Gametocytes

For live-cell fluorescence imaging, samples of mature gametocyte culture (> 0.3% exflagellation) were stained with 500 nM SiR-tubulin (Spirochrome) for 3 hours at 37°C. Samples were additionally stained with 5 μg/ml wheat germ agglutinin (WGA) conjugated to AlexaFluor488 (Invitrogen) and 500 nM Vybrant^™^ DyeCycle^™^ Violet for 30 minutes at 37°C. Stained samples were protected from light and strictly maintained at 37°C, to prevent premature activation, until imaging.

Samples were treated with 10 μM 1294^46^,10 μM ML10^35^, 25 μM Bortezomib or DMSO and normalised to 0.25% DMSO for 3 hours at 37°C, before imaging.

### Imaging Live Microgametogenesis by Widefield Microscopy

Wells of an Ibidi 8-well μ-Slide were pre-treated with 140 μl ookinete medium and slides were pre-positioned on the microscope stage. To activate gametogenesis, 6 μL of the stained gametocytes, equating to ~30 million total erythrocytes, was added directly to ookinete media-treated wells at room temperature (21°C). Samples were imaged with a Prime 95B sCMOS camera (photometrics) on a Nikon Ti2-E widefield microscope using x 100 Plan Apo 1.4 numerical aperture (NA) oil objective with NIS Elements v4.20 software. SiR-Tubulin, WGA-488 and Vybrant^™^ DyeCycle^™^ Violet staining were imaged with a Cy5, GFP and DAPI filter set. The triggered multi-wavelength LED and static quad band filter cube was used in acquisition through Z and between wavelengths.

Early stages of microgametogenesis (0-10 minutes) were acquired as 3D datasets through time at ‘no delay’ and with 34 ms exposure time. Z-stacks were acquired at 0.2 μm steps from above and below the cell with a Piezo driven stage. Acquisition was alternated between brightfield and fluorescence to minimise phototoxicity and subsequently prolong parasite viability. Exflagellation was captured as 2D (single z-slice) datasets, acquiring with no delay between frames.

### Image Analysis

The open-source bioimage analysis software Icy^47^ was used to analyse all time-lapse datasets. All 3D datasets are depicted here as 2D maximum intensity projections. Early developmental stages of microgametogenesis, acquired through t and z, were deconvolved using a custom-made Protocol in Icy. The Protocol is attached in the **Additional Supplementary Information**. Within the Protocol, a Sequence File Batch loop locates the 3D+t files from which a channel of interest is extracted. The selected channel of each time frame is processed as an individual 3D stack. The metadata of each stack is read and used as an input for the EpiDEMIC deconvolution bloc (Epifluorescence Deconvolution MICroscopy), a blind (i.e. without Point Spread Function (PSF) knowledge) deconvolution method for widefield fluorescence microscopy 3D data^48^. All timelapses were deconvolved over 50 iterations and 2 loops.

All 2D timelapse data and 3D data depicted as 2D maximum intensity projections or 3D sectioned views were created using Icy. To quantify SiR-tubulin and Vybrant^™^ DyeCycle^™^ Violet intensity, egress phenotypes and cell circularity, raw 3D data was converted to 2D maximum intensity projections in NIS Elements v4.20 prior to analysis. Egress phenotypes were quantified by manual observation. To measure the circularity and staining intensity of individual cells, each cell was defined as a custom region of interest. The SiR-tubulin and Vybrant^™^ DyeCycle^™^ Violet staining intensity of each individual cell was quantified using the time-measurement function in NIS Elements v4.20 and is reported in arbitrary units. The circularity of each cell was measured using the Automated Measurement feature in NIS Elements v4.20. Circularity is reported here as shape measure values from 0-1 derived from the area and perimeter of each cell, with higher values representing shapes of increasing circularity and circles being characterised as a value of 1. All graphical and statistical data was analysed with GraphPad Prism version 8.0.

## Supporting information

Supplementary Videos and Protocol

## ACKNOWLEDGEMENTS

We thank Irene Garcia-Barbazan, Eliana Real and David Grimson for assisting with parasite culture and for sharing expert transmission advice. Additional thanks to other members of the Baum lab for helpful discussions and experimental support, with particular thanks to Thomas C. A. Blake for assistance in data production. We also thank Robert E. Sinden for valuable discussions about the biology of microgametogenesis. Finally, we thank collaborators in the Baker lab and Van Voorhis lab for their generous donation of microgametogenesis inhibitors 1294 and ML10, respectively.

## AUTHOR CONTRIBUTIONS

S.Y., S.J., G.A. and J.B. designed experiments. S.Y., S.J., H.S., D.G. and G.A. performed and analysed experiments. S.Y., S.J., M.T.F, A.C, F.D and E.R. cultured *P. falciparum*. D.C.A.G designed the Icy batch deconvolution protocol.

## COMPETING INTERESTS

The authors declare no competing interests.

## FUNDING

S.Y. is supported by a Ph.D. studentship from an EPSRC Doctoral Training Partnership award (Grant EP/R512540/1) to Imperial College London. JB was supported by an Investigator Award from Wellcome (100993/Z/13/Z). This work was funded in part by an award to JB from the Bill and Melinda Gates Foundation (OPP1181199). The Facility for Imaging by Light Microscopy (FILM) at Imperial College London is part-supported by funding from the Wellcome Trust (grant 104931/Z/14/Z) and BBSRC (grant BB/L015129/1)

## Notes

### Competing Interest Statement

The authors have declared no competing interest.

## REFERENCES

1. Josling, G.A. & Llinás, M. Sexual development in Plasmodium parasites: knowing when it’s time to commit. Nat Rev Microbiol 13, 573–87 (2015).

2. Dixon, M.W., Dearnley, M.K., Hanssen, E., Gilberger, T. & Tilley, L. Shape-shifting gametocytes: how and why does P. falciparum go banana-shaped? Trends Parasitol 28, 471–8 (2012).

3. Dixon, M.W.A. & Tilley, L. Plasmodium falciparum goes bananas for sex. Mol Biochem Parasitol 244, 111385 (2021).

4. Billker, O., Shaw, M.K., Margos, G. & Sinden, R.E. The roles of temperature, pH and mosquito factors as triggers of male and female gametogenesis of Plasmodium berghei in vitro. Parasitology 115 (Pt 1), 1–7 (1997).

5. Andreadaki, M. et al. Sequential Membrane Rupture and Vesiculation during Plasmodium berghei Gametocyte Egress from the Red Blood Cell. Scientific Reports 8, 3543 (2018).

6. Guttery, D.S., Roques, M., Holder, A.A. & Tewari, R. Commit and Transmit: Molecular Players in Plasmodium Sexual Development and Zygote Differentiation. Trends in Parasitology 31, 676–685 (2015).

7. Andreadaki, M. et al. Sequential Membrane Rupture and Vesiculation during Plasmodium berghei Gametocyte Egress from the Red Blood Cell. Sci Rep 8, 3543 (2018).

8. Ponzi, M. et al. Egress of Plasmodium berghei gametes from their host erythrocyte is mediated by the MDV-1/PEG3 protein. Cell Microbiol 11, 1272–88 (2009).

9. Talman, A.M. et al. PbGEST mediates malaria transmission to both mosquito and vertebrate host. Mol Microbiol 82, 462–74 (2011).

10. Deligianni, E. et al. Critical role for a stage-specific actin in male exflagellation of the malaria parasite. Cell Microbiol 13, 1714–30 (2011).

11. Deligianni, E. et al. A perforin-like protein mediates disruption of the erythrocyte membrane during egress of Plasmodium berghei male gametocytes. Cell Microbiol 15, 1438–55 (2013).

12. Olivieri, A. et al. Distinct properties of the egress-related osmiophilic bodies in male and female gametocytes of the rodent malaria parasite Plasmodium berghei. Cell Microbiol 17, 355–68 (2015).

13. Sinden, R.E., Canning, E.U., Bray, R.S. & Smalley, M.E. Gametocyte and gamete development in Plasmodium falciparum. Proc R Soc Lond B Biol Sci 201, 375–99 (1978).

14. Billker, O. et al. Azadirachtin disrupts formation of organised microtubule arrays during microgametogenesis of Plasmodium berghei. J Eukaryot Microbiol 49, 489–97 (2002).

15. Sinden, R.E., Canning, E.U. & Spain, B. Gametogenesis and fertilization in Plasmodium yoelii nigeriensis: a transmission electron microscope study. Proc R Soc Lond B Biol Sci 193, 55–76 (1976).

16. Sinden, R.E., Talman, A., Marques, S.R., Wass, M.N. & Sternberg, M.J. The flagellum in malarial parasites. Curr Opin Microbiol 13, 491–500 (2010).

17. Sinden, R.E. Malaria, sexual development and transmission: retrospect and prospect. Parasitology 136, 1427–34 (2009).

18. Bell, A. Microtubule Inhibitors as Potential Antimalarial Agents. Parasitology Today 14, 234–240 (1998).

19. Rawlings, D.J. et al. α-Tubulin II is a male-specific protein in Plasmodium falciparum. Molecular and Biochemical Parasitology 56, 239–250 (1992).

20. Angrisano, F., Tan, Y.H., Sturm, A., McFadden, G.I. & Baum, J. Malaria parasite colonisation of the mosquito midgut--placing the Plasmodium ookinete centre stage. Int J Parasitol 42, 519–27 (2012).

21. Ruecker, A. et al. A male and female gametocyte functional viability assay to identify biologically relevant malaria transmission-blocking drugs. Antimicrob Agents Chemother 58, 7292–302 (2014).

22. Bertiaux, E. et al. Expansion microscopy provides new insights into the cytoskeleton of malaria parasites including the conservation of a conoid. PLoS Biol 19, e3001020 (2021).

23. Sinden, R.E. Gametocytogenesis of Plasmodium falciparum in vitro: an electron microscopic study. Parasitology 84, 1–11 (1982).

24. Ashdown, G.W. et al. A machine learning approach to define antimalarial drug action from heterogeneous cell-based screens. bioRxiv (2019).

25. Delves, M.J. et al. A high throughput screen for next-generation leads targeting malaria parasite transmission. Nat Commun 9, 3805 (2018).

26. Geoghegan, N.D. et al. 4D analysis of malaria parasite invasion offers insights into erythrocyte membrane remodeling and parasitophorous vacuole formation. Nature Communications 12, 3620 (2021).

27. Zeeshan, M. et al. Protein phosphatase 1 regulates atypical mitotic and meiotic division in Plasmodium sexual stages. Communications Biology 4, 760 (2021).

28. Wall, R.J. et al. Plasmodium APC3 mediates chromosome condensation and cytokinesis during atypical mitosis in male gametogenesis. Sci Rep 8, 5610 (2018).

29. Lukinavičius, G. et al. Fluorogenic probes for live-cell imaging of the cytoskeleton. Nature Methods 11, 731–733 (2014).

30. Dubois, J. et al. Fluorescent and biotinylated analogues of docetaxel: Synthesis and biological evaluation. Bioorganic & Medicinal Chemistry 3, 1357–1368 (1995).

31. Fang, H. et al. Multiple short windows of calcium-dependent protein kinase 4 activity coordinate distinct cell cycle events during Plasmodium gametogenesis. eLife 6, e26524 (2017).

32. Hitz, E., Balestra, A.C., Brochet, M. & Voss, T.S. PfMAP-2 is essential for male gametogenesis in the malaria parasite Plasmodium falciparum. Scientific Reports 10, 11930 (2020).

33. Delves, M.J. et al. Routine in vitro culture of P. falciparum gametocytes to evaluate novel transmission-blocking interventions. Nat Protoc 11, 1668–80 (2016).

34. Doggett, J.S., Ojo, K.K., Fan, E., Maly, D.J. & Van Voorhis, W.C. Bumped kinase inhibitor 1294 treats established Toxoplasma gondii infection. Antimicrob Agents Chemother 58, 3547–9 (2014).

35. Baker, D.A. et al. A potent series targeting the malarial cGMP-dependent protein kinase clears infection and blocks transmission. Nat Commun 8, 430 (2017).

36. McRobert, L. et al. Gametogenesis in Malaria Parasites Is Mediated by the cGMP-Dependent Protein Kinase. PLOS Biology 6, e139 (2008).

37. Bennink, S., Kiesow, M.J. & Pradel, G. The development of malaria parasites in the mosquito midgut. Cellular Microbiology 18, 905–918 (2016).

38. Yahiya, S. et al. Plasmodium falciparum protein Pfs16 is a target for transmission-blocking antimalarial drug development. bioRxiv, 2021.06.14.448287 (2021).

39. McRobert, L. et al. Gametogenesis in malaria parasites is mediated by the cGMP-dependent protein kinase. PLoS Biol 6, e139 (2008).

40. Reynolds, J.M., El Bissati, K., Brandenburg, J., Günzl, A. & Mamoun, C.B. Antimalarial activity of the anticancer and proteasome inhibitor bortezomib and its analog ZL3B. BMC clinical pharmacology 7, 13–13 (2007).

41. Adams, J. et al. Proteasome Inhibitors: A Novel Class of Potent and Effective Antitumor Agents. Cancer Research 59, 2615–2622 (1999).

42. Chen, D., Frezza, M., Schmitt, S., Kanwar, J. & Dou, Q.P. Bortezomib as the first proteasome inhibitor anticancer drug: current status and future perspectives. Curr Cancer Drug Targets 11, 239–53 (2011).

43. Fu, X. et al. Ubiquitin-Like Proteasome System Represents a Eukaryotic-Like Pathway for Targeted Proteolysis in Archaea. mBio 7(2016).

44. Tarrason Risa, G. et al. The proteasome controls ESCRT-III–mediated cell division in an archaeon. Science 369, eaaz2532 (2020).

45. Xie, S.C. et al. Target Validation and Identification of Novel Boronate Inhibitors of the Plasmodium falciparum Proteasome. J Med Chem 61, 10053–10066 (2018).

46. Ojo, K.K. et al. A specific inhibitor of PfCDPK4 blocks malaria transmission: chemical-genetic validation. J Infect Dis 209, 275–84 (2014).

47. de Chaumont, F. et al. Icy: an open bioimage informatics platform for extended reproducible research. Nat Methods 9, 690–6 (2012).

48. Soulez, F., Denis, L., Tourneur, Y. & Thiébaut, É. Blind deconvolution of 3D data in wide field fluorescence microscopy. in International Symposium on Biomedical Imaging CDROM (2012).

