## Supplementary Information for "4D live-cell imaging of microgametogenesis in the human malaria parasite *Plasmodium falciparum*"

**Key words:**

**SUPPLEMENTARY FIGURES**

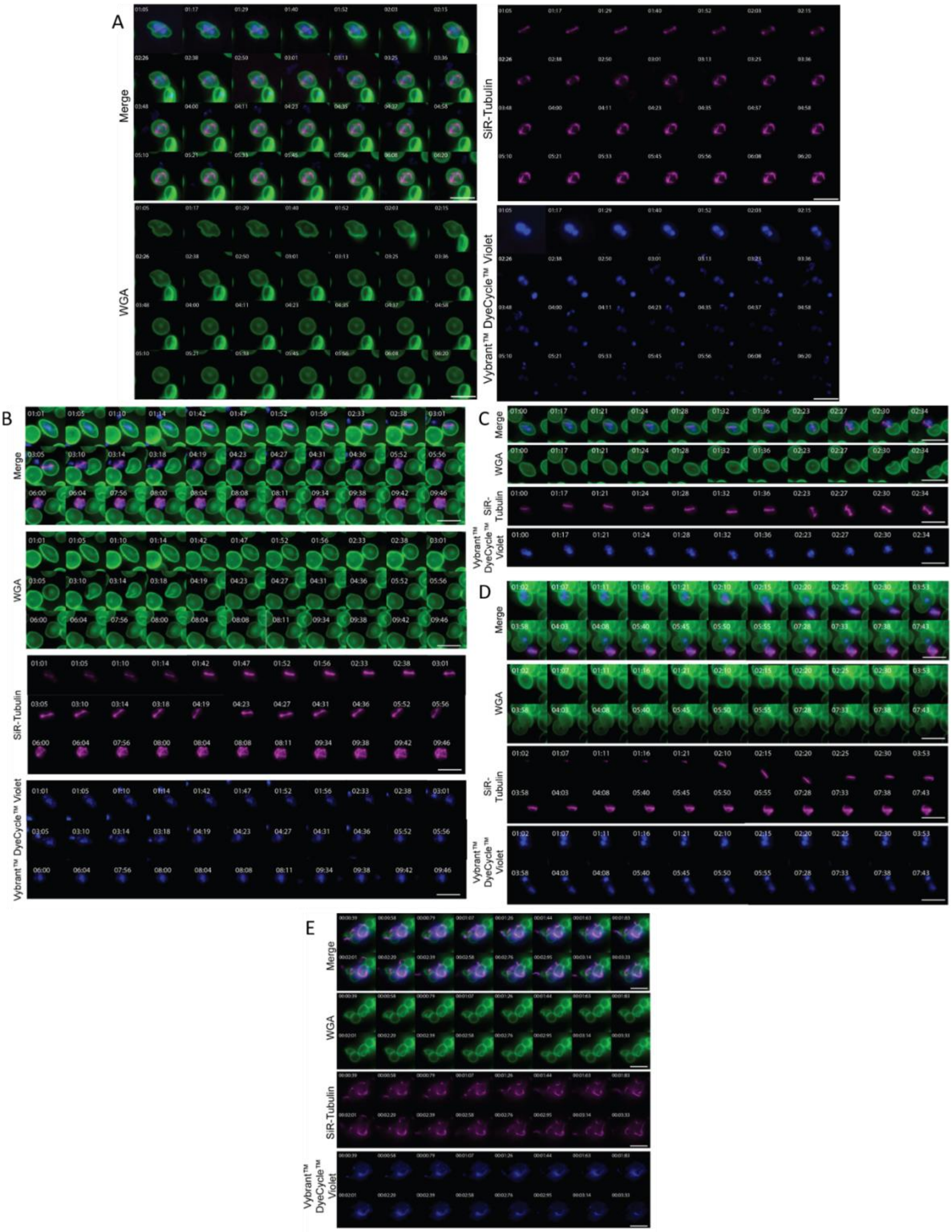


**Figure S1. Individual channels of 3-colour *P. falciparum* microgametogenesis timelapse data**

Individual channels of **(A)** tubulin dynamics from **Figure 2B**, **(B)** host erythrocyte egress from **Figure 2D, (C-D)** additional egress data and **(E)** exflagellation from **Figure 3C**. Merged channels, microtubules (SiR-Tubulin), host erythrocyte membrane (WGA) and parasite nuclei (Vybrant™ DyeCycle™ Violet) of 2D maximum intensity projection data is shown. Time is depicted as minutes and seconds (mm:ss) in **A-D** and minutes, seconds and milliseconds (mm:ss:ms) in **E**. Scale bars = 10 µm.

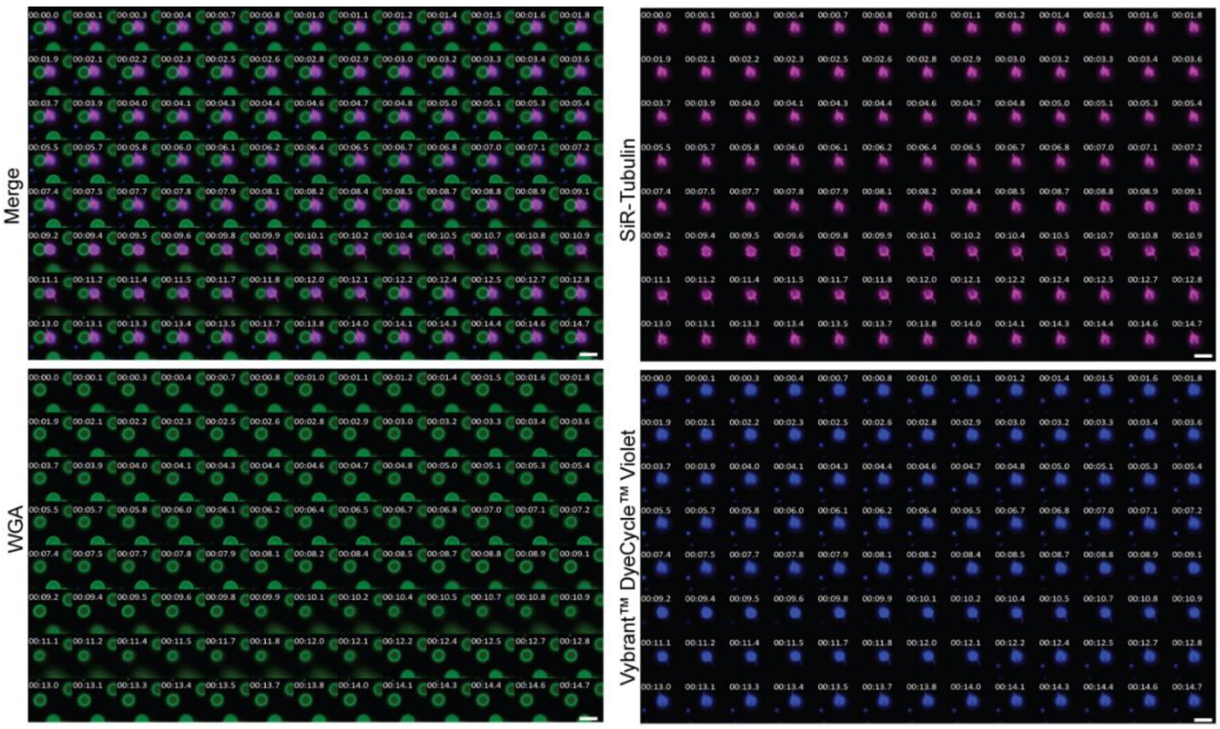


**Figure S2. Early emergence of *P. falciparum* microgametes**

The merged image, microtubules (SiR-Tubulin), host erythrocyte membrane (WGA) and parasite nuclei (Vybrant™ DyeCycle™ Violet) of microgametogenesis in the early stages of exflagellation. Images represent stills derived from timelapses, portrayed as 2D maximum intensity projection of 3D data. See **Supplementary Video 7** for the corresponding timelapse. Time is depicted as minutes, seconds and milliseconds (mm:ss:ms). Scale bars = 10 µm.

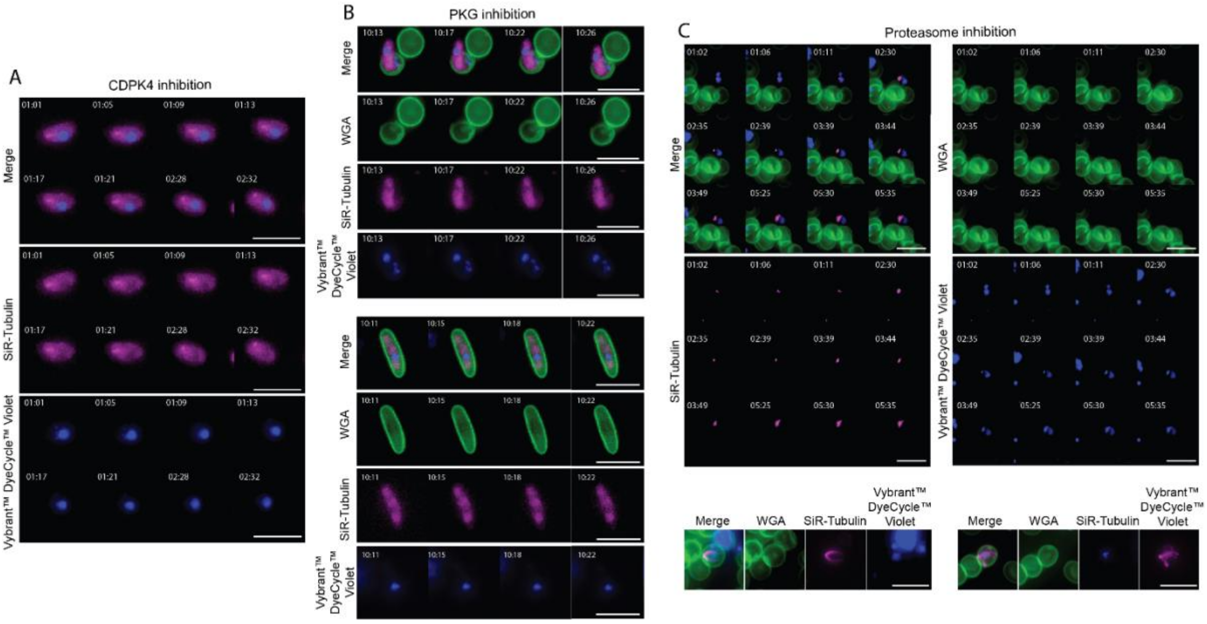


**Figure S3. Individual channels of CDPK4, PKG and proteasome-inhibited parasites**

Individual channels depicting the phenotypes of **(A)** CDPK4, **(B)** PKG and **(C)** proteasome-inhibition by 1294, ML10 and bortezomib, respectively. 2D maximum intensity projection images are depicted and accompany **Figures 4A-C**. Merged channels, microtubules (SiR-Tubulin), host erythrocyte membrane (WGA) and parasite nuclei (Vybrant™ DyeCycle™ Violet) are shown. Time is depicted as minutes and seconds (mm:ss). Scale bars = 10 µm.

### DESCRIPTION OF ADDITIONAL SUPPLEMENTARY FILES

File: **Supplementary file 1. Batch Deconvolution Protocol.protocol**

Description: Custom-made protocol file comprised of processing steps, termed Blocs, for batch deconvolution of 3D timelapse data in Icy.

File: **Supplementary video 1**

Description: SiR-tubulin (magenta) stained developing microgametocytes during microgametogenesis. The microtubule organising centre (MTOC) within the falciform gametocyte is shown to transform, a spindle forms and axonemes are nucleated and elongated from basal bodies. Accompaniment for **Figure 2A**.

File: **Supplementary video 2**

Description: 3D sectioned view of early developmental stages of microgametogenesis. SiR-Tubulin (magenta), WGA-488 (green) and Vybrant™ DyeCycle™ Violet (blue). The 2D maximum intensity projection (MIP) of 3D frames is shown. Accompaniment for **Figure 2B** and **S1A**.

File: **Supplementary video 3**

Description: 3D sectioned view of early microgametogenesis and egress. SiR-Tubulin (magenta), WGA-488 (green) and Vybrant™ DyeCycle™ Violet (blue). The 2D maximum intensity projection (MIP) of 3D frames is shown. Accompaniment for **Figure 2C** and **S1B**.

File: **Supplementary video 4**

Description: Maximum intensity projection of early microgametogenesis, from mitotic spindle to lengthened axonemes. SiR-Tubulin (magenta), WGA-488 (green) and Vybrant™ DyeCycle™ Violet (blue). The 2D maximum intensity projection (MIP) of 3D frames is shown.

File: **Supplementary video 5**

Description: 3D sectioned view of early microgametogenesis and egress. SiR-Tubulin (magenta), WGA-488 (green) and Vybrant™ DyeCycle™ Violet (blue). The 2D maximum intensity projection (MIP) of 3D frames is also shown. Accompaniment for **Figure S1C**.

File: **Supplementary video 6**

Description: 3D sectioned view of early microgametogenesis and egress. SiR-Tubulin (magenta), WGA-488 (green) and Vybrant™ DyeCycle™ Violet (blue). The 2D maximum intensity projection (MIP) of 3D frames is also shown. Accompaniment for **Figure S1D**.

File: **Supplementary video 7**

Description: 2D maximum intensity projection (MIP) of 3D timelapse data depicting early microgamete emergence. **(A)** SiR-Tubulin (magenta), WGA-488 (green) and Vybrant™ DyeCycle™ Violet (blue) (accompaniment for **Figure S2**) and corresponding **(B)** brightfield frames.

File: **Supplementary video 8**

Description: 2D (single Z-slice) timelapse of exflagellation, brightfield information depicted. Accompaniment for **Figure 3A**.

File: **Supplementary video 9**

Description: 2D (single Z-slice) timelapse of exflagellation, brightfield information depicted.

File: **Supplementary video 10**

Description: 2D (single Z-slice) timelapse of exflagellation of a parasite stained with SiR-tubulin (magenta). Accompaniment for **Figure 3B**.

File: **Supplementary video 11**

Description: 2D (single Z-slice) timelapse of exflagellation a parasite stained with SiR-tubulin (magenta).

File: **Supplementary video 12**

Description: 2D (single Z-slice) timelapse of exflagellation of a parasite stained with SiR-tubulin (magenta).

File: **Supplementary video 13**

Description: 2D (single Z-slice) timelapse of exflagellation of a parasite stained with SiR-tubulin (magenta).

File: **Supplementary video 14**

Description: 2D (single Z-slice) timelapse of exflagellation of a parasite stained with SiR-tubulin (magenta).

File: **Supplementary video 15**

Description: 2D (single Z-slice) timelapse of exflagellation of a parasite stained with SiR-tubulin (magenta), WGA-488 (green) and Vybrant™ DyeCycle™ Violet (blue). Accompaniment for **Figure 3C** and **S1E**.

File: **Supplementary video 16**

Description: 2D (single Z-slice) timelapse of exflagellation of a parasite stained with SiR-tubulin (magenta), WGA-488 (green) and Vybrant™ DyeCycle™ Violet (blue).

File: **Supplementary video 17**

Description: Rotated view of 3D view of an exflagellating parasite stained with SiR-tubulin (magenta), WGA-488 (green) and Vybrant™ DyeCycle™ Violet (blue). A single time-frame is shown. Accompaniment for **Figure 3E**.

File: **Supplementary video 18**

Description: 2D maximum intensity projection (MIP) of 3D timelapse data depicting **1294**-treatment phenotype. SiR-Tubulin (magenta), WGA-488 (green) and Vybrant™ DyeCycle™ Violet (blue).

File: **Supplementary video 19**

Description: 3D sectioned view of **1294**-treatment phenotype. SiR-Tubulin (magenta), WGA-488 (green) and Vybrant™ DyeCycle™ Violet (blue). The 2D maximum intensity projection (MIP) of 3D frames is shown. Accompaniment for **Figure 4A**.

File: **Supplementary video 20**

Description: 2D maximum intensity projection (MIP) of 3D timelapse data depicting **1294**-treatment phenotype. SiR-Tubulin (magenta), WGA-488 (green) and Vybrant™ DyeCycle™ Violet (blue).

File: **Supplementary video 21**

Description: 2D maximum intensity projection (MIP) of 3D timelapse data depicting **1294**-treatment phenotype. SiR-Tubulin (magenta), WGA-488 (green) and Vybrant™ DyeCycle™ Violet (blue).

File: **Supplementary video 22**

Description: 2D maximum intensity projection (MIP) of 3D timelapse data depicting **1294**-treatment phenotype. SiR-Tubulin (magenta), WGA-488 (green) and Vybrant™ DyeCycle™ Violet (blue).

File: **Supplementary video 23**

Description: 3D sectioned view of **ML10**-treatment phenotype. SiR-Tubulin (magenta), WGA-488 (green) and Vybrant™ DyeCycle™ Violet (blue). The 2D maximum intensity projection (MIP) of 3D frames is shown. Accompaniment for **Figure 4B**.

File: **Supplementary video 24**

Description: 3D sectioned view of **ML10**-treatment phenotype. SiR-Tubulin (magenta), WGA-488 (green) and Vybrant™ DyeCycle™ Violet (blue). The 2D maximum intensity projection (MIP) of 3D frames is shown. Accompaniment for **Figure 4B**.

File: **Supplementary video 25**

Description: 3D sectioned view of **Bortezomib**-treatment phenotype. SiR-Tubulin (magenta), WGA-488 (green) and Vybrant™ DyeCycle™ Violet (blue). The 2D maximum intensity projection (MIP) of 3D frames is shown. Accompaniment for **Figure 4C**.

File: **Supplementary video 26**

Description: 2D maximum intensity projection (MIP) of 3D timelapse data depicting **Bortezomib**-treatment phenotype. SiR-Tubulin (magenta), WGA-488 (green) and Vybrant™ DyeCycle™ Violet (blue).

File: **Supplementary video 27**

Description: 2D maximum intensity projection (MIP) of 3D timelapse data depicting **Bortezomib**-treatment phenotype. Accompaniment for **Figure 4C**.

File: **Supplementary video 28**

Description: 2D maximum intensity projection (MIP) of 3D timelapse data depicting **Bortezomib**-treatment phenotype. Accompaniment for **Figure 4C**.
